# Metapipeline-DNA: A Comprehensive Germline & Somatic Genomics Nextflow Pipeline

**DOI:** 10.1101/2024.09.04.611267

**Authors:** Yash Patel, Chenghao Zhu, Takafumi N. Yamaguchi, Nicholas K. Wang, Nicholas Wiltsie, Nicole Zeltser, Alfredo E. Gonzalez, Helena K. Winata, Yu Pan, Mohammed Faizal Eeman Mootor, Timothy Sanders, Sorel T. Fitz-Gibbon, Cyriac Kandoth, Julie Livingstone, Lydia Y. Liu, Benjamin Carlin, Aaron Holmes, Jieun Oh, John Sahrmann, Shu Tao, Stefan Eng, Rupert Hugh-White, Kiarod Pashminehazar, Andrew Park, Arpi Beshlikyan, Madison Jordan, Selina Wu, Mao Tian, Jaron Arbet, Beth Neilsen, Roni Haas, Yuan Zhe Bugh, Gina Kim, Joseph Salmingo, Wenshu Zhang, Aakarsh Anand, Edward Hwang, Anna Neiman-Golden, Philippa Steinberg, Wenyan Zhao, Prateek Anand, Raag Agrawal, Brandon L. Tsai, Paul C. Boutros

**Author notes:** These authors contributed equally to this work (Yash Patel, Chenghao Zhu, Takafumi N. Yamaguchi). Corresponding author: Dr. Paul C. Boutros, University of California Los Angeles, Los Angeles, California, 90095.

## Abstract

**Summary:** The price, quality and throughout of DNA sequencing continue to improve. Algorithmic innovations have allowed inference of a growing range of features from DNA sequencing data, quantifying nuclear, mitochondrial and evolutionary aspects of both germline and somatic genomes. To automate analyses of the full range of genomic characteristics, we created an extensible Nextflow meta-pipeline called metapipeline-DNA. Metapipeline-DNA analyzes targeted and whole-genome sequencing data from raw reads through pre-processing, feature detection by multiple algorithms, quality-control and data-visualization. Each step can be run independently and is supported robust software engineering including automated failure-recovery, robust testing and consistent verifications of inputs, outputs and parameters. Metapipeline-DNA is cloud-compatible and highly configurable, with options to subset and optimize each analysis. Metapipeline-DNA facilitates high-scale, comprehensive analysis of DNA sequencing data.

**Availability:** Metapipeline-DNA is an open-source Nextflow pipeline under the GPLv2 license and is available at https://github.com/uclahs-cds/metapipeline-DNA.

## Introduction

High-throughput technologies have made biomedical research increasingly data-intensive. DNA sequencing is a key enabling technology, used both in routine clinical care and to support a wide range of research studies^1^. Ongoing improvements in DNA sequencing continue to reduce costs and enable new discoveries, like elucidation complex structural variants (SVs) and repetitive genomic regions by long-read sequencing^2^. Modern germline DNA sequencing studies routinely quantify single-nucleotide polymorphisms (SNPs), SVs, telomere length, mitochondrial copy number and variation, copy number and many other features^3–5^.

DNA sequencing has been especially helpful in characterizing tumors. Cancers are characterized by widespread genomic rearrangements, variation in mutation clonality, specific patterns of somatic mutations associated with carcinogens or other features and a host of features absent or uncommon in germline genomes like kataegis and chromothripsis^6^. Comprehensive analyses of cancer sequencing can improve diagnosis, prognosis and management^7,8^. In many studies both a sample of a cancer and a “reference” normal sample from the same individual are sequenced to better distinguish somatic from germline variation and enable analysis of germline-somatic interactions.

The growing availability of DNA sequencing has been paralleled by rapid development and adoption of both specific algorithms and workflow software. New discoveries often rely heavily on complex workflows comprising a mixture of established and novel algorithms^9^. These workflows, often termed “pipelines”, are implemented in a range of orchestration frameworks including Galaxy^10^, Snakemake^11^, Common Workflow Language (CWL)^12^ and Nextflow^13^. Workflows provide a way to automate processes by minimizing manual handling of data flow and facilitating stitching together of different tools to process raw data into refined forms such as lists of variants or quantitation of specific features.

The use of complex workflows has placed a growing emphasis on standardization, extensibility, quality control and compute infrastructure. Workflow implementations routinely differ across research groups, with many groups creating their own. Many workflows lack key features like unit testing, integration testing, error-handling, fault-tolerance, input-output verification, quality-control, data-visualization and use of multiple algorithms to create consensus calls^14^. Given the volume of data and the expense of compute, workflows are often bespoke to the high-performance computing environment used by a single group^15^. Portability of workflows to new environments is part of the “model to data” (M2D) paradigm in data sharing and processing^16^. M2D overcomes the cost, time and privacy risks of data-transfer by bringing models or algorithms to the computing system where data is stored. M2D thus necessitates that models be portable across providers and environments to support workflow usage in conjunction with good data management principles hinging on findability, accessibility, interoperability and reusability^17^.

To address the need for a robust open-source DNA sequencing analysis pipeline, we created metapipeline-DNA. This Nextflow meta-pipeline is highly customizable and is capable of processing data from any stage of analysis. It can process DNA sequencing data starting from raw reads through alignment and recalibration, variant calling and even highly integrated analyses likely tumour subclonal reconstruction. Extensive quality control, testing and data-visualization are built into each individual step and into the full metapipeline. It can work on multiple compute systems and clouds, facilitating analyses at any scale.

## Results

### Overview

metapipeline-DNA is a Nextflow meta-pipeline for analysis of DNA sequencing data. It can analyze both targeted and whole-genome sequencing with 16 pipelines (**Table 1**) that collectively transform raw sequencing reads into sets of detected variants and other genetic and evolutionary features (**Figure 1A**). Most individual pipelines can execute multiple alternative algorithms and create consensus calls. For example, subclonal copy number aberration detection uses two algorithms (FACETS and Battenberg) and produces visualizations including logR and BAF plots (**Figure 1B**). Similarly four separate algorithms can be executed for somatic single nucleotide variant (SNV) detection^14^, automatically generating a consensus set of predictions and variant-associated data-visualizations (**Figure 1C-D**). Each pipeline can be executed independently and can be extensively parameterized to customize the selection and tuning of algorithms.

**Table 1:**
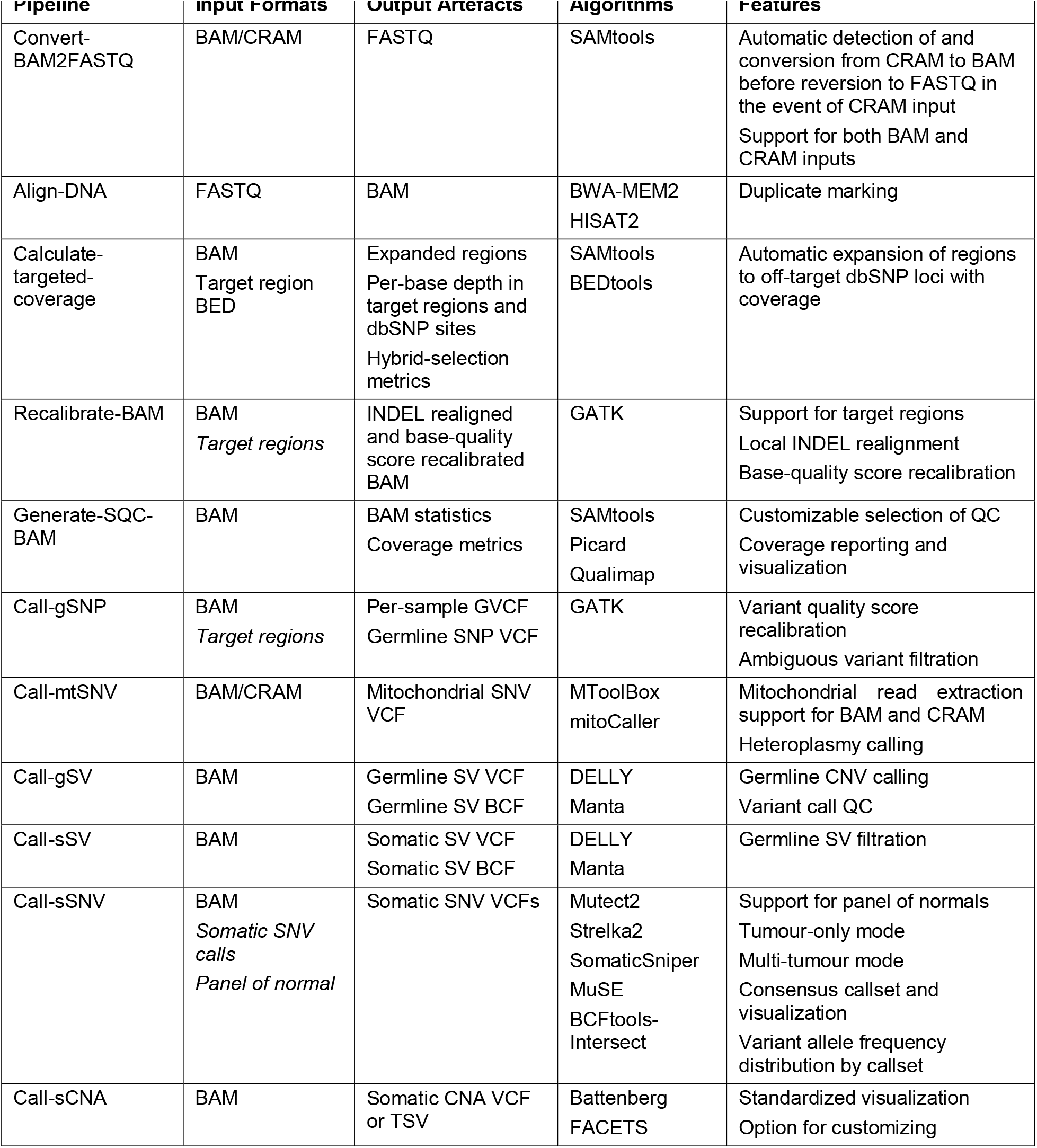
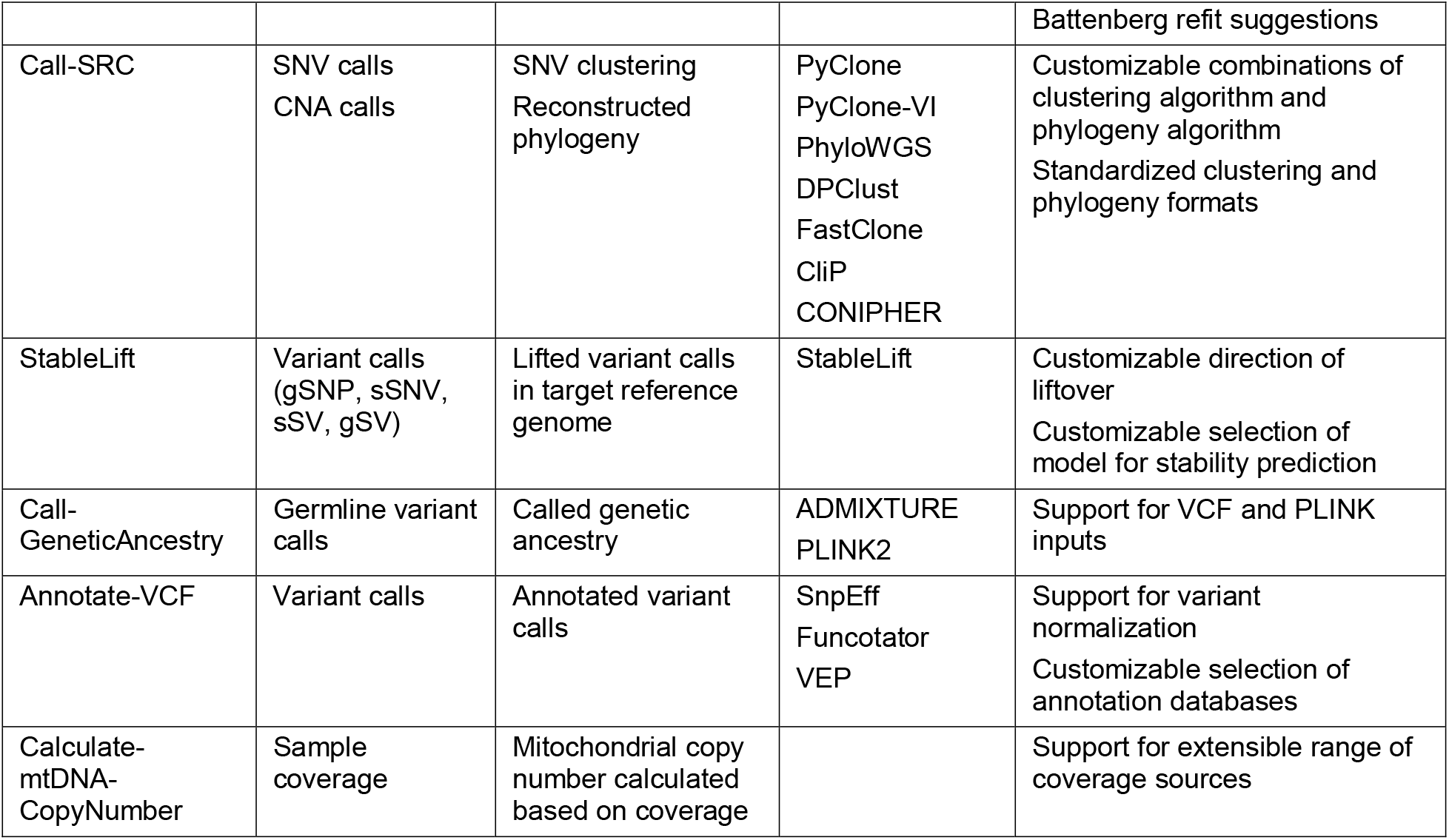
metapipeline-DNA Constituent Pipelines. Pipelines encompassed within metapipeline-DNA and their inputs, outputs, algorithms, and key features. Inputs that are *italicized* are optional and inputs separated by “/” represent a list of choices from which one must be chosen.

**Figure 1:**
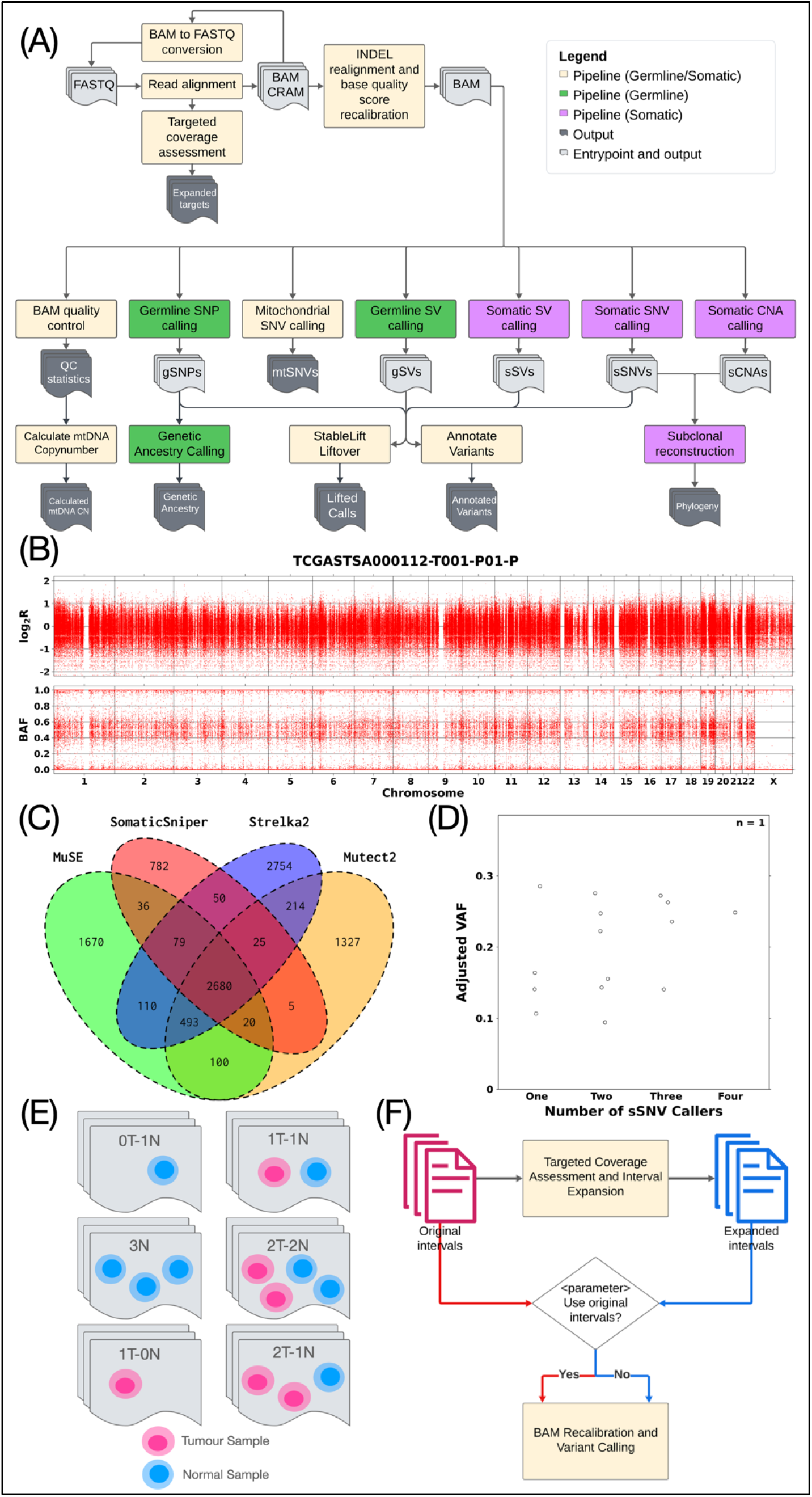
Data flow and visualizations. **A**. Data flow through metapipeline-DNA. **B**. Normalized tumour coverage relative to the matched normal (log_2_R) and the B-allele frequency of individual SNPs laid out across the genome to support CNA detection. **C**. Example intersection diagram of consensus variants between 4 SNV callers: MuSE2, SomaticSniper, Strelka2 and Mutect2. **D**. Variant allele frequencies based on consensus between callers. VAFs are indicated for all combinations of consensus between one, two, three and four variant callers, with each data point representing one combination. The adjusted VAF is calculated as an average of all variants present in the combination. **E**. Sample combinations supported by metapipeline-DNA. The nT-mN (ex. 2T-2N) combination indicates any arbitrary numbers of normal and tumour samples. Each combination is automatically detected and considered during processing of all pipelines to select appropriate algorithms and processing modalities. **F**. Automatic and customizable interval usage in metapipeline-DNA. Original intervals undergo assessment and expansion at dbSNP sites with high coverage to produce expanded intervals. The option of using the expanded or the original intervals for downstream BAM recalibration and variant calling is parameterized with both options automated.

Several different sample run-modes are available, which we denote with the terminology *n*T-*m*N, where *n* indicates the number of tumour samples and *m* the number of reference samples (**Figure 1E**). Thus classic paired tumour-normal analysis is 1T-1N. Metapipeline-DNA fully supports modes like 0T-1N (*i*.*e*. germline DNA sequencing), 0T-3N (*e*.*g*. family trios), 1T-0N (*i*.*e*. unpaired tumour-only sequencing) and arbitrary multi-region tumour sequencing (*e*.*g*. 5T-1N). The primary limitation to multi-sample analyses is compute resource availability – particularly RAM and scratch-disk space. Metapipeline-DNA automatically handles input types for each mode and only executes feasible pipelines, independent of user-selections. For example, in 0T modes, variant detection is restricted to germline variants without users having to provide manual guidance or parameterization.

The default mode of metapipeline-DNA accepts unaligned reads in FASTQ^18^ format and executes all pipelines. A range of alternative entry-points are possible, including starting from an unaligned BAM, an aligned BAM^19^ or from CRAM files, with automatic BAM-to-FASTQ conversions as needed. A few pipelines accept alternative entry-points, such as SNV and copy number aberration (CNA) calls for tumour subclonal reconstruction^20^ (**Figure 1A**). Documentation of all dependencies, input and output formats is available on standardized structured GitHub pages: current states at writing are summarized in **Supplementary Table 1**.

We engineered metapipeline-DNA to be intrinsically flexible with all necessary dependencies automatically identified and executed based on user selection. All run-modes and dependency identification have defaults set to the most common behaviour across thousands of runs, but with easy parameterization. For example, when input data is already aligned the default is to use these alignments. Configuration parameters allow the user to control whether reads are converted to FASTQ and re-aligned and whether aligned reads are recalibrated and so forth.

In a similar way, metapipeline-DNA is flexible to the specific genome build used, and has been tested extensively with GRCh37, GRCh38 and GRCm39. It can run in two modes: WGS mode and targeted-sequencing mode, based on user parameterization. Targeted-sequencing mode supports all subsets of the genome, including exome sequencing and arbitrary panels. Options are available to assess coverage, expand targets with off-target coverage sites and automatically use expanded target intervals for downstream processing (**Figure 1F**).

## Data Visualization & Quality-Control

metapipeline-DNA includes a range of quality control steps and pipelines to assess data quality at many levels, including reads, alignments and variant calls. The pipeline for back-conversion from BAM/CRAM to FASTQ includes built-in checks, including SAM flag and alignment statistics generation and read count comparison before and after conversion to FASTQ (to ensure no loss of reads due to file corruption or parallelization scatter-gather failures, for example). These quality-controls produce a variety of data-visualizations and reports. For example, alignment quality is inferred from BAM (or CRAM) files in a range of ways including coverage distributions over the genome (or target region with or without padding; **Figure 2A-B**). Reads are quantified by a range of quality metrics, including total counts, mapping qualities, GC content, insert sizes, read lengths, duplications and others. **Figure 2C** shows an example of read number stratified by a range of quality groupings. A range of software are used to generate these metrics, including SAMtools^19^, Picard^21^ and Qualimap^22^. Pileup summaries at common sites are generated and used as a precursor to estimate contamination across samples. Visualization is also built into the SV calling pipelines to produce representations of structural variants and their categorization (inversion, insertion, breakend/translocation) in circos plots (**Figure 2D**).

**Figure 2:**
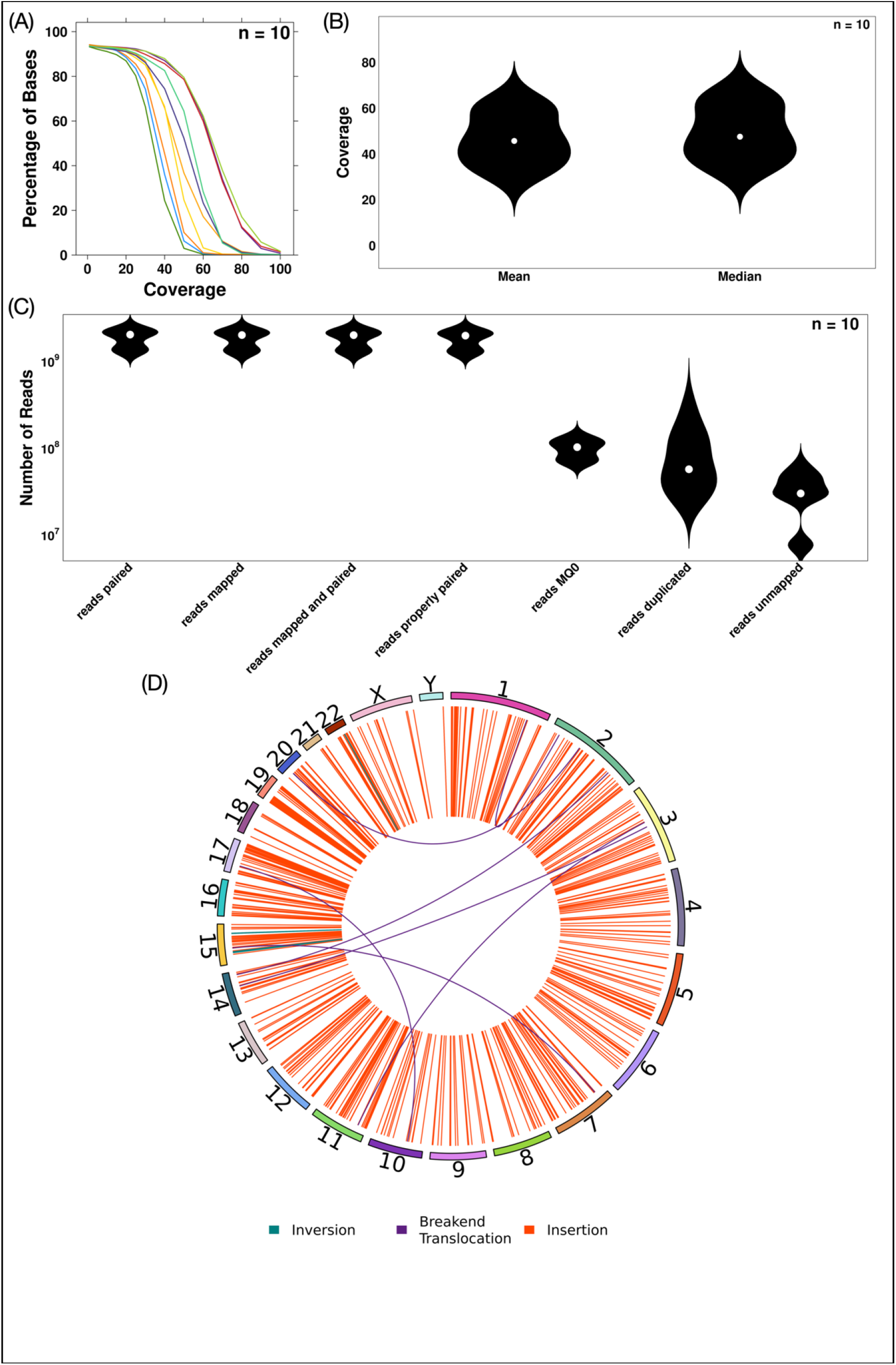
Alignment and coverage metrics. **A**. Percent of bases in the genome at each fold of coverage for normal and tumour samples (each line represents one sample) for all five WGS PCAWG patients. Each line represents a different sample, with the percentage of bases calculated using the coverage metrics. **B**. Distribution of mean and median coverage across all samples, highlighting two rough separations arising from the normal samples and tumour samples with the normal samples being in the lower coverage separation. **C**. Distributions of read numbers across alignment metrics including mapped/unmapped, low mapping quality, duplication and paired. **D**. Circos plot of somatic structural variants categorized into inversions, insertions and breakend/translocations.

In targeted-sequencing mode, additional coverage assessment is performed through per-base read depth calculations at target regions and well-characterized off-target polymorphic sites provided from dbSNP^23^. The workflow also generates an expanded set of targets encompassing the original target regions plus user-defined polymorphic sites (typically dbSNP) enriched in coverage over a user-defined threshold. Metapipeline-DNA provides with configuration to automatically use the expanded targets with BAM recalibration and variant calling pipelines.

Variants are additionally evaluated for stability across reference genomes: StableLift^24^ is available as an optional workflow to support liftover of sSVs, gSVs, sSNVs and gSNPs between GRCh37 and GRCh38. In addition to liftover, the pipeline annotates variants with databases such as dbSNP^23^ and applies a model to assign a stability score to each variant to indicate the likelihood of the variant being consistently represented across the two reference genome builds. Variant type-specific assessment is also performed. For example, germline SNP calls undergo filtration using models built from variant quality scores for both SNPs and Indels. Somatic SNVs are assessed based on consensus between callers and associated variant allele frequencies. The consensus approach across callers allows for filtering of SNV calls to reduce the rate of false positives made by a single caller.

Germline SNP calls undergo genetic sex-specific evaluations to reduce the rate of false positives. Variants on chromosomes X and Y in non-pseudo-autosomal regions^25–27^ (PARs) are extracted and filtered based on the genetic sex. In XY samples, heterozygous genotype calls are removed and homozygous genotype calls are converted to hemizygous. In XX samples, all chromosome Y variant calls are removed. This reduces the false positive rate of variant calls made on the sex chromosomes.

### Software-Engineering & Pipeline Robustness

We placed a heavy focus on generating re-usable and extensible software that could automatically detect and recover from common errors, particularly in the compute environment. This led us to adopt or create a series of development practices and pipeline features aimed at maximizing quality. All software is open-source, available on GitHub (https://github.com/uclahs-cds/metapipeline-DNA), with transparent tracking of issues and discussions. Development followed a test-driven approach using NFTest^14^. Metapipeline-DNA has a suite of 95 unit, integration and regression tests that are run for each new release with testing performed for different stages of execution from end-to-end tests to individual pipeline tests. The tests utilize publicly available simulated sequencing data from the ICGC-TCGA DREAM Somatic Mutation Calling Tumour Heterogeneity (SMC-Het) Challenge^28^ sub-sampled at various sequencing depths to facilitate different tests. Our extensive use of Docker containers allows seamless co-existence of multiple pipeline versions, and the combination of automated testing and containerization facilitates rapid updating with new features or dependency versions. Standardized GitHub issue templates support robust reporting of both bugs and new feature-requests, allowing ideal collaboration (**Supplementary Figure 1**). At writing, development has involved 43 contributors making 1,382 pull-requests and 46 individuals making 1,117 suggestions, feature-requests and issue-reports across 17 repositories.

Bioinformatics data has high intrinsic variability, and bioinformatics software can be prone to significant numbers of failures – particularly in heterogeneous computing environments. Failure handling is built into metapipeline-DNA to predict and minimize wasted computation. We automated input and parameter validation to catch issues prior to commitment of compute resources^29^. Proactive validation of pipeline parameters is implemented to avoid errors prior to resource commitment. Individual pipelines are modularized and fault-tolerant such that errors or failures in one pipeline stay isolated from and do not terminate other pipelines that are not direct dependencies. metapipeline-DNA can be easily re-run in cases of failure, triggered starting from prior partial results with a simple parameterization.

All outputs are organized with standardized directory and naming structures (**Supplementary Figure 2**). Filenames have been standardized to provide dataset, organism and sample information in a consistent way across pipelines. metapipeline-DNA similarly organizes log-files to ensure saving of and ready access to the metapipeline-DNA logs, individual pipeline-level logs and compute partition logs. These logs capture execution and resource usage metrics for every process. Robust tooling has been developed around process and pipeline execution to ensure logs are captured for both successful and failing steps to enable debugging and record-keeping. Scripts have been created that automatically “crawl” over a series of pipeline runs to extract and tabulate information about run success, compute resources and other features.

### Compute Infrastructure

metapipeline-DNA includes compute-agnostic customizability of execution and scheduling in distributed workflows. It has been tested and validated on both the Azure and AWS clouds. Execution follows the pattern of a single leading job responsible for submission and monitoring of per-sample or per-patient analysis jobs. Execution is currently performed with the Slurm executor with optional specification of compute partitions^30^. Parameters also exist to control rate of job submission and amount of parallelization/resources usage. Once configured and submitted, metapipeline-DNA automatically handles processing of an entire cohort with input parsing and job submission without user intervention. Real-time monitoring is available through email notifications sent from a server watching individual step start, end and status. The choice of executor itself is parameterized and can be easily extended to other environments.

metapipeline-DNA includes optimizations for disk usage, including (optional) eager intermediate file removal and built in checks to allow for optimized disk usage (performing I/O operations from high-performance working disks). Resource allocation for individual steps is also automatically handled, with pipelines running in parallel as available resources allows. Resource-related robustness is also built into pipelines to detect memory allocation failures from individual tools and automatically retry processes with higher allocations.

### Case Studies

We assessed the performance of germline SNP XY filtration using the Genome in a Bottle (GIAB) HG002 sample^31^. Variant calls generated by pipeline-call-gSNP were assessed through comparison with the GIAB HG002 XY small variant benchmark v1.0 as the truth set^31^. Both raw variant calls and XY filtered variant calls were compared to demonstrate a decrease in false positive (FP) variant calls from 1,347 INDELs (FDR = 0.054) to 221 INDELs (FDR = 0.009) and 7,563 SNPs (FDR = 0.083) to 1,290 SNPs (FDR = 0.015). The true positive (TP) and false negative (FN) rates remain consistent through filtration, demonstrating the effectiveness of XY filtration in improving precision without adversely affecting sensitivity. **Figure 3A** shows these results, highlighting elimination of 1,126 false-positive INDEL calls and 6,273 false-positive SNP calls in a single sample from this procedure.

**Figure 3:**
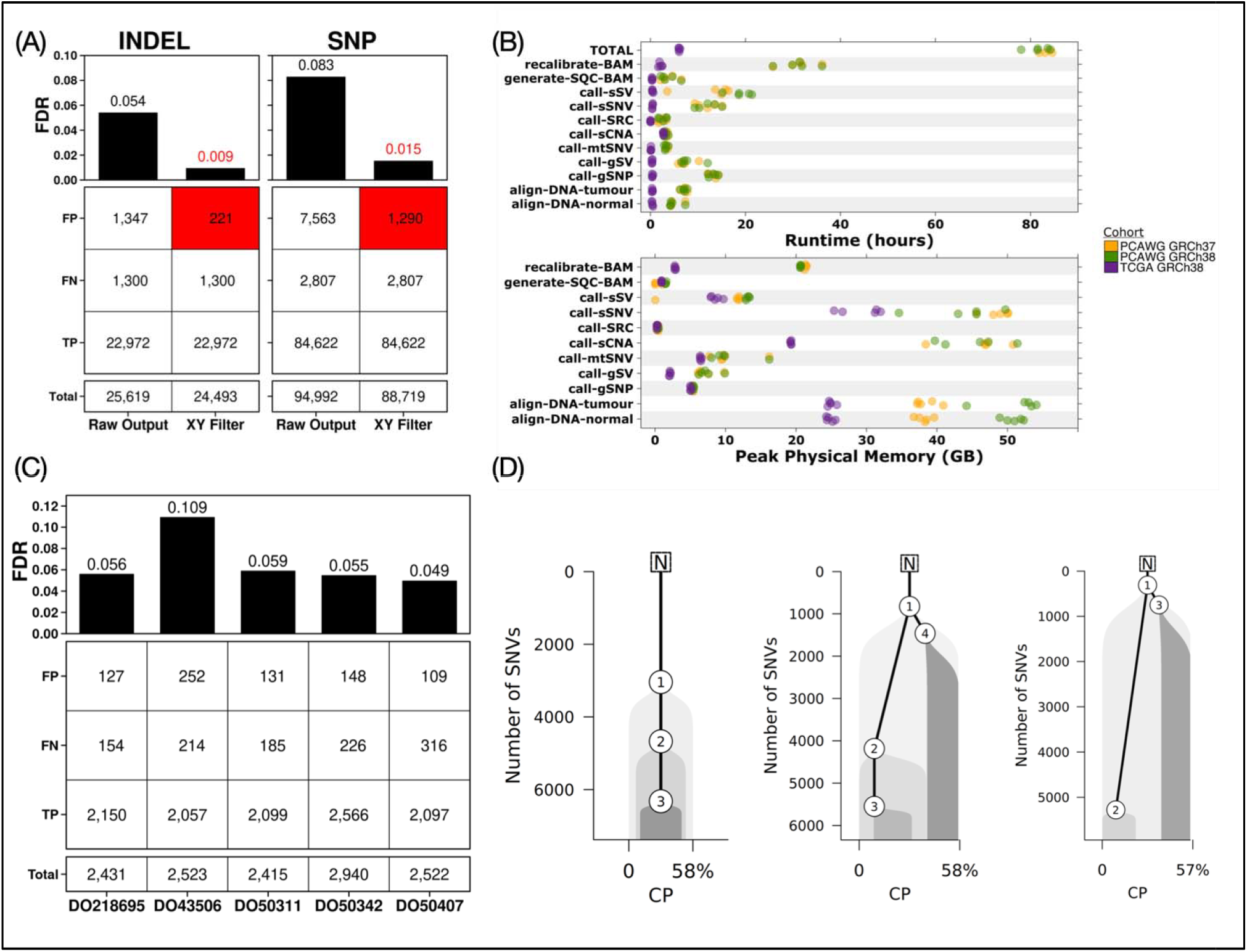
Use-cases and benchmarking. **A**. TP, FN and FP variant calls comparing raw SNP and INDEL calls with XY filtered variant calls against the GIAB HG002 truth set. Numbers represent the number of variant calls with reduction in false discovery rate with filtration highlighted in red. **B**. Time and memory usage of pipelines per sample for the three different processing cohorts: PCAWG with GRCh38, PCAWG with GRCh37 and TCGA with GRCh38. Time is measured as wall-clock time taken by each pipeline and the total time taken by metapipeline-DNA. Memory is measured as the peak RAM usage by any single process by any pipeline. **C**. TP, FN and FP variant calls comparing consensus call-SNV calls from the PCAWG-5 samples against a set of validation variant calls made from targeted deep-sequencing of the same samples. Numbers represent the number of variant calls. **D**. Reconstructed phylogeny of tumour samples SA478344, SA528788 and SA528876 using consensus SNV callset comprising variants called by at least two out of four SNV callers (MuSE2, SomaticSniper, Strelka2, Mutect2) and FACETS CNAs. Nodes represent identified subclones with the evolutionary history depicted over SNV accumulation. Along the x-axis is the cellular prevalence (CP), indicating the fraction of all cells comprising each subclone.

As an additional demonstration and benchmark, ten normal-tumour pairs were processed through the entirety of metapipeline-DNA. Five pairs were selected from the Pan-Cancer Analysis of Whole Genomes (PCAWG)^32^ 63 dataset and another five from The Cancer Genome Atlas (TCGA)^33^. The PCAWG-63 samples were sequenced with whole-genome sequencing and derived from multiple cancer types: one from uterine corpus endometrial carcinoma, one from biliary tract carcinoma, and three from esophageal adenocarcinoma. The samples had a median coverage of 63x (range: 45-65x) for the tumour samples and 38x (range: 34-54x) for the normal samples. The TCGA samples were derived from soft tissue sarcoma samples sequenced with exome-targeted sequencing. Both pairs were processed using metapipeline-DNA from alignment to subclonal reconstruction. The PCAWG-63 samples were processed with both GRCh38 and GRCh37, with similar runtimes across the two reference builds at an average of 81.76 hours (95% CI: ± 14.23) for GRCh38 and 83.36 hours (95% CI: ± 12.99) for GRCh37. Across the ten pairs, memory usage peaked in call-sSNV (average ± 95% CI: 48.54GB ± 2.30 and 29.32GB ± 3.82 for PCAWG63 GRCh37 and TCGA GRCh38 respectively) and in align-DNA (average ± 95% CI: 51.42 ± 5.07 for PCAWG63 GRCh38). Runtimes and peak memory usage of metapipeline-DNA for these samples are visualized in **Figure 3B** and summarized in **Supplementary Table 2**. Both run-times and memory usage are a function of compute hardware and parameter selection and can be extensively tuned and optimized.

We assessed our consensus-based sSNV calling workflow and its consensus callset using the PCAWG samples with targeted deep-sequencing (mean coverage of 653x) validation as a truth set. The true positive rate (TPR) for the samples ranged from 0.87 to 0.93 with false discovery rate (FDR) ranging from 0.05 to 0.11 (**Figure 3C**). To demonstrate phylogenetic reconstruction, we subsampled 5,000 SNVs and used CNA calls (average 138 per sample; **Figure 3D**). Variant allele frequencies aggregated over all combinations of consensus calls are shown for all samples in **Supplementary Figure 3**.

## Discussion

metapipeline-DNA was designed to facilitate analysis of DNA sequencing data at scale while retaining the configurability and flexibility needed in academic environments. This is a key contrast to field programmable gate array (FPGA) approaches such as DRAGEN^34^, which attain outstanding speed through hardware optimization at the expense of algorithmic flexibility and evolution. As the field of genomics evolves, the ability to quickly integrate and test emerging methods continues to be extremely important, highlighting a limitation of fixed-function hardware solutions.

metapipeline-DNA fills this key niche of supporting the rapidly expanding volume of sequencing data, supporting a range of existing tools and algorithms and remaining flexible for ongoing expansion. By easing and optimizing the multi-step analyses intrinsic to DNA sequencing data, it reduces the barrier to incorporating new methods and analyzing large datasets. Indeed, it is entirely feasible for metapipeline-DNA to leverage and incorporate FPGA-enabled and graphics processing units (GPU)-accelerated methods directly as part of its modular structure (*e*.*g*. for alignment); this is a key area of ongoing development.

Individual pipelines within metapipeline-DNA are modular, creating a plug-and-play architecture that can be adapted to support additional technologies as they become available. Algorithms and workflows for processing long-read data, for example, pose an avenue for expanding the meta-pipeline as such tools mature and long-read datasets become more common. The context of DNA also brings up the possibility of similar meta- pipelines for other biological molecules such as RNA and proteins. Workflows across different biomolecules can share the architecture, automation and quality-control of metapipeline-DNA in a way that allows improvements to any single pipeline to improve the others. Such workflows are currently under development to provide a similar level of configurability and extensibility for analyses of RNA and protein data.

The volume of data and size of individual samples being generated and processed in sequencing studies is often very large. With that comes a need for optimization of analysis pipelines’ data handling. Metapipeline-DNA contains several disk usage optimizations to efficiently handle large amounts of data while minimizing I/O operations and cross-file system data movement. The framework connecting analyses automatically identifies necessary outputs from dependent pipelines and makes it available without any redundant copying or duplication. There are additional enhancements that are underway to minimize duplicated data and disk usage of metapipeline-DNA by building plugins to enable moving of files rather than copying when possible and optimizing individual pipelines to avoid shuffling around large output files.

metapipeline-DNA is a highly customizable DNA sequencing analysis pipeline combining speed and flexibility in a modular framework to enable processing of data at any point from read alignment to tumour subclonal reconstruction. By facilitating the integration of diverse tools and supporting the rapid development of new methodologies, it positions itself as a versatile platform for future enhancements as novel DNA sequencing and analysis methods are developed.

## Methods

### Analysis Cohort

To demonstrate the use of metapipeline-DNA, we chose ten normal-tumour pairs. Five were WGS pairs from PCAWG-63: one from uterine corpus endometrial carcinoma donor DO43506, one from biliary tract carcinoma donor DO218695 and three from esophageal adenocarcinoma donors DO50342, DO50407 and DO50311. Five were exome sequencing pairs of soft tissue sarcoma pairs from TCGA donors TCGA-QQ-A8VD, TCGA-X6-A8C6, TCGA-HS-A5N8, TCGA-DX-A1L2 and TCGA-HB-A2OT^32,33^.

### Alignment and Variant Calling

Sequencing reads were aligned to the GRCh38.p7 reference build including decoy contigs from GATK using BWA-MEM2^35^ (v2.2.1) in paired-end mode followed by duplicate marking with MarkDuplicatesSpark using GATK^36^ (v4.2.4.1). For the GRCh37 runtime benchmarking, alignment was performed to the GRCh37 reference build including decoy contigs. The results alignments were recalibrated through Indel realignment using GATK (v3.7.0) and base-quality score recalibration using GATK (v4.2.4.1). Quality metrics were generated using SAMtools^19^ (v1.18) stats and Picard^21^ (v3.1.0) CollectWgsMetrics. Germline SNPs were called using HaplotypeCaller from GATK (v4.2.4.1) followed by variant recalibration using GATK (v4.2.4.1). Germline SNPs underwent XY filtration using Hail^37^ (v0.2.113) with benchmarking assessment performed using Hap.py^38^ (v0.3.15). Genetic ancestry was called using germline SNPs and INDELs as inputs using ADMIXTURE^39^ (v1.3.0) and PLINK^40,41^ (2.00a4.5lm). Germline SVs were called using Delly2^42^ (v1.2.6) and Manta^43^ (v1.6.0). Mitochondrial SNVs were called using mitoCaller^3^ (v1.0.0). Somatic SNVs were called using MuSE2^44^ (v2.0.4), SomaticSniper^45^ (v1.0.5.0), Strelka2^46^ (v2.9.10) and Mutect2^36^ (v4.5.0.0) followed by a consensus workflow to identify variants called by two or more callers using BCFtools^47^ (v1.17) with quality control plots generated with BPG^48^ (v7.1.0) and VennDiagram^49^ (v1.7.4). Somatic SVs were called using Delly2^42^ (v1.2.6) and Manta^43^ (v1.6.0) and visualized with circlize^50^ (v0.4.16). Somatic CNAs were called using CNV_FACETS^51^ (v0.16.0) for the PCAWG sample and using Battenberg^52^ (v2.2.9) for the TCGA sample with visualization generated using BPG^48^ (v7.1.0). Taking the consensus set of somatic SNV calls and the CNA calls, subclonal reconstruction was performed using PyClone-VI^53^ (v0.1.2), PhyloWGS^54^ (v2205be1) and FastClone^55^ (v1.0.9). Variant liftover was performed using BCFtools^47^ (v1.20) and stability prediction using StableLift^24^ (v1.0.0). Variant annotation was done using SnpEff^56^ (v5.1d), Funcotator^36^ (v4.2.4.1) and VEP^57^ (v101.0) and ClinVar^58^ (v20211016). Reconstructed phylogeny was visualized using CEV^59^ (v2.0.0). Data validation was performed with PipeVal^29^ (v5.1.0) and data processing was done using Nextflow^13^ (v23.04.2).

## Acknowledgements

The authors gratefully acknowledge the ongoing support of all present and past members of the Boutros lab in providing suggestions, practical use-cases and support. The authors also acknowledge the Office of Health Informatics and Analytics at UCLA Health IT for their infrastructure support, particularly high-performance compute provisioning, data management and resource tuning.

