## Supplementary Table for "Metapipeline-DNA: A Comprehensive Germline & Somatic Genomics Nextflow Pipeline"

| Pipeline | Input Data | Sample Modes | Output Artefacts | Algorithms |
|---|---|---|---|---|
| Convert-BAM2FASTQ (https://github.com/uclahs-cds/pipeline-convert-BAM2FASTQ) | BAM/CRAM – Aligned reads in BAM or CRAM format | Single sample | FASTQ – Raw reads extracted per readgroup | SAMtools v1.15.1 |
| Align-DNA (https://github.com/uclahs-cds/pipeline-align-DNA) | FASTQ – Paired raw reads with information about sequencing such as readgroup, library, sequencing center | Single sample | BAM – Aligned reads in BAM format | BWA-MEM2 v2.2.1<br>HISAT2 v2.2.1 |
| Calculate-targeted-coverage (https://github.com/uclahs-cds/pipeline-calculate-targeted-coverage) | BAM – Aligned reads in BAM format<br>Target region BED – Genomic sites targeted for sequencing | Single sample | Expanded regions<br>Per-base depth in target regions and dbSNP sites<br>Hybrid-selection metrics | SAMtools v1.16.1<br>BEDtools v2.29.2 |
| Recalibrate-BAM (https://github.com/uclahs-cds/pipeline-recalibrate-BAM) | BAM – Aligned reads in BAM format<br>*Target regions – Genomic sites targeted for sequencing/analysis* | Single sample<br>Normal-tumour paired samples<br>Multi-normal and/or multi-tumour samples | INDEL realigned and base-quality score recalibrated BAM | GATK v3.7.0, v4.2.4.1 |
| Generate-SQC-BAM (https://github.com/uclahs-cds/pipeline-generate-SQC-BAM) | BAM – Aligned reads in BAM format (typically including INDEL realignment and BQSR) | Single sample | BAM statistics – Statistics related to alignment, reads, quality, duplication<br>Coverage metrics – Statistics and plots of coverage across the genome | SAMtools v1.18<br>Picard v3.1.0<br>Qualimap v2.3 |
| Call-gSNP (https://github.com/uclahs-cds/pipeline-call-gSNP) | BAM - Aligned reads in BAM format (typically including INDEL realignment and BQSR)<br>*Target regions - Genomic sites targeted for sequencing/analysis* | Single sample<br>Normal-tumour paired samples<br>Multi-normal and/or multi-tumour samples | Per-sample GVCF – Genomic VCF generated per sample<br>Germline SNP VCF – Recalibrated and filtered germline SNP calls for set of given samples | GATK v4.2.4.1 |
| Call-mtSNV (https://github.com/uclahs-cds/pipeline-call-mtSNV) | BAM/CRAM – Aligned reads in BAM or CRAM format (typically including INDEL realignment and BQSR) | Single sample<br>Norma-tumour paired samples | Mitochondrial SNV VCF | MToolBox v1.2.1-b52269e<br>mitoCaller v1.0.0 |
| Call-gSV (https://github.com/uclahs-cds/pipeline-call-gSV) | BAM – Aligned reads in BAM format (typically including INDEL realignment and BQSR); single normal BAM | Single sample | Germline SV BCF – Variant calls made by DELLY in BCF format<br>Germline SV BCF – Variant calls made by Manta in VCF format | DELLY v1.2.6<br>Manta v1.6.0 |
| Call-sSV https://github.com/uclahs-cds/pipeline-call-sSV) | BAM – Aligned reads in BAM format | Normal-tumour paired samples | Somatic SV BCF – Variant calls made | DELLY v1.2.6<br>Manta v1.6.0 |

| | | | | |
|---|---|---|---|---|
| | (typically including INDEL realignment and BQSR) | | by DELLY in BCF format<br>Somatic SV VCF – Variant calls made by Manta in VCF format | |
| Call-sSNV<br>https://github.com/uclahs-cds/pipeline-call-sSNV) | BAM – Aligned reads in BAM format (typically including INDEL realignment and BQSR)<br>*Somatic SNV calls – Variant calls provided in VCF format to run the consensus call workflow*<br>*Panel of normal – PON used with Mutect2 to improve variant calls* | Single tumour sample<br>Normal-tumour paired samples<br>Multiple tumour samples | Somatic SNV VCFs – Variant calls made by each of the algorithms, VCFs separated per algorithm and per variant type (SNV, MNV, INDEL) when applicable | Mutect2 v4.5.0.0<br>Strelka2 v2.9.10<br>SomaticSniper v1.0.5.0<br>MuSE v2.0.4<br>BCFtools-Intersect v1.17 |
| Call-sCNA<br>(https://github.com/uclahs-cds/pipeline-call-sCNA) | BAM – Aligned reads in BAM format (typically including INDEL realignment and BQSR) | Normal-tumour paired samples | Somatic CNA TSV – Aberrations called by Battenberg in TSV format<br>Somatic CNA VCF – Aberrations called by FACETS in VCF format | Battenberg v2.2.9<br>FACETS v0.16.0 |
| Call-SRC<br>(https://github.com/uclahs-cds/pipeline-call-SRC) | SNV calls – Generated by any of the algorithms from call-sSNV<br>CNA calls – Generated by any of the algorithms from call-sCNA and HATCHet | Single tumour sample<br>Multiple tumour samples | SNV clustering – Result of clustering of SNVs by clustering algorithms<br>Reconstructed phylogeny | PyClone v0.13.1<br>PyClone-VI v0.1.2<br>PhyloWGS v2205be1<br>DPClust v75f5d7e<br>FastClone v1.0.9<br>CliP v1.3<br>CONIPHER v2.2.0 |
| StableLift<br>(https://github.com/uclahs-cds/pipeline-StableLift) | Variant calls – Generated by any of the following algorithms: HaplotypeCaller, Mutect2, Strelka2, SomaticSniper, MuSE2, DELLY2 | Single sample | Lifted variant calls – Variant calls lifted over into the target reference genome<br>Variant stability score – Predicted score of variant stability across reference genome builds | BCFtools v1.20<br>StableLift v1.0.0 |
| Call-GeneticAncestry<br>(https://github.com/uclahs-cds/pipeline-call-GeneticAncestry) | Germline variant calls – Generated by any germline variant caller | Cohort of samples | Predicted genetic ancestry | ADMIXTURE v1.3.0<br>PLINK2 v2.00a4.5lm |
| Annotate-VCF<br>(https://github.com/uclahs-cds/pipeline-annotate-VCF) | Variant calls – Generated by any caller in VCF format | Single sample | Annotated variant calls – Variant calls annotated with the selected databases | SnpEff v5.1d<br>Funcotator v4.2.4.1<br>VEP v101.0 |
| Calculate-mtDNA-CopyNumber<br>(https://github.com/uclahs- | Genomic coverage – Coverage information per-contig to be used | Single sample | Calculated mitochondrial DNA copy number | |

| | |
|---|---|
| cds/pipeline-calculate-mtDNA-CopyNumber) | in calculating mitochondrial DNA copy number |

**Supplementary Table 1: Detailed pipeline inputs, outputs, and tools.** Detailed description of inputs, outputs, run modes, and tools encompassed in metapipeline-DNA. Inputs that are *italicized* are optional and inputs separated by "/" represent a list of choices from which one must be chosen.

| Pipeline | PCAWG WGS GRCh37 (wall-clock time in hours) | PCAWG WGS GRCh37 (Peak RAM in GB) | PCAWG WGS GRCh38 (wall-clock time in hours) | PCAWG WGS GRCh38 (Peak RAM in GB) | TCGA WXS GRCh38 (wall-clock time in hours) | TCGA WXS GRCh38 (Peak RAM in GB) |
|---|---|---|---|---|---|---|
| Align-DNA (normal) | 4.92 ± 1.68 | 38.12 ± 1.36 | 5.56 ± 2.04 | 50.82 ± 1.73 | 0.40 ± 0.12 | 24.82 ± 0.69 |
| Align-DNA (tumour) | 7.06 ± 0.69 | 38.48 ± 1.98 | 8.20 ± 0.86 | 51.42 ± 5.07 | 0.39 ± 0.09 | 24.96 ± 0.62 |
| Recalibrate-BAM | 31.01 ± 4.64 | 21.20 ± 0.44 | 31.31 ± 3.08 | 20.62 ± 0.06 | 2.10 ± 0.44 | 2.84 ± 0.07 |
| Generate-SQC-BAM | 3.80 ± 2.17 | 0.41 ± 0.66 | 4.96 ± 0.67 | 1.53 ± 0.08 | 0.34 ± 0.07 | 0.92 ± 0.009 |
| Call-gSNP | 13.14 ± 1.16 | 5.36 ± 0.07 | 7.27 ± 3.41 | 5.42 ± 0.06 | 0.41 ± 0.11 | 5.06 ± 0.07 |
| Call-mtSNV | 3.43 ± 0.17 | 10.56 ± 4.06 | 3.23 ± 0.42 | 10.58 ± 4.01 | 0.096 ± 0.024 | 6.46 ± 0.11 |
| Call-sSNV | 12.04 ± 2.92 | 48.54 ± 2.30 | 10.52 ± 1.83 | 43.7 ± 6.98 | 0.43 ± 0.07 | 29.32 ± 3.82 |
| Call-sSV | 12.85 ± 6.60 | 9.47 ± 6.55 | 18.89 ± 3.01 | 13.14 ± 0.29 | 0.44 ± 0.11 | 8.60 ± 0.91 |
| Call-gSV | 7.33 ± 2.09 | 7.24 ± 1.90 | 8.05 ± 2.83 | 7.48 ± 1.80 | 0.34 ± 0.08 | 2.10 ± 0.09 |
| Call-sCNA | 3.43 ± 0.17 | 46.0 ± 5.67 | 3.23 ± 0.42 | 45.14 ± 5.89 | 2.72 ± 0.05 | 19.26 ± 0.07 |
| Call-SRC | 2.39 ± 0.96 | 0.41 ± 0.09 | 2.58 ± 1.01 | 0.41 ± 0.09 | 0.014 ± 0.002 | 0.27 ± 0.04 |
| **TOTAL** | **83.36 ± 12.99** | **-** | **81.76 ± 14.23** | **-** | **6.05 ± 0.80** | **-** |

**Supplementary Table 2: Runtime and peak physical memory usage of pipelines per sample with 95% confidence intervals.** The total runtime is less than the sum of the individual pipelines' runtimes due to parallelization of variant calling pipelines.
