## Supplementary Figure for "Metapipeline-DNA: A Comprehensive Germline & Somatic Genomics Nextflow Pipeline"

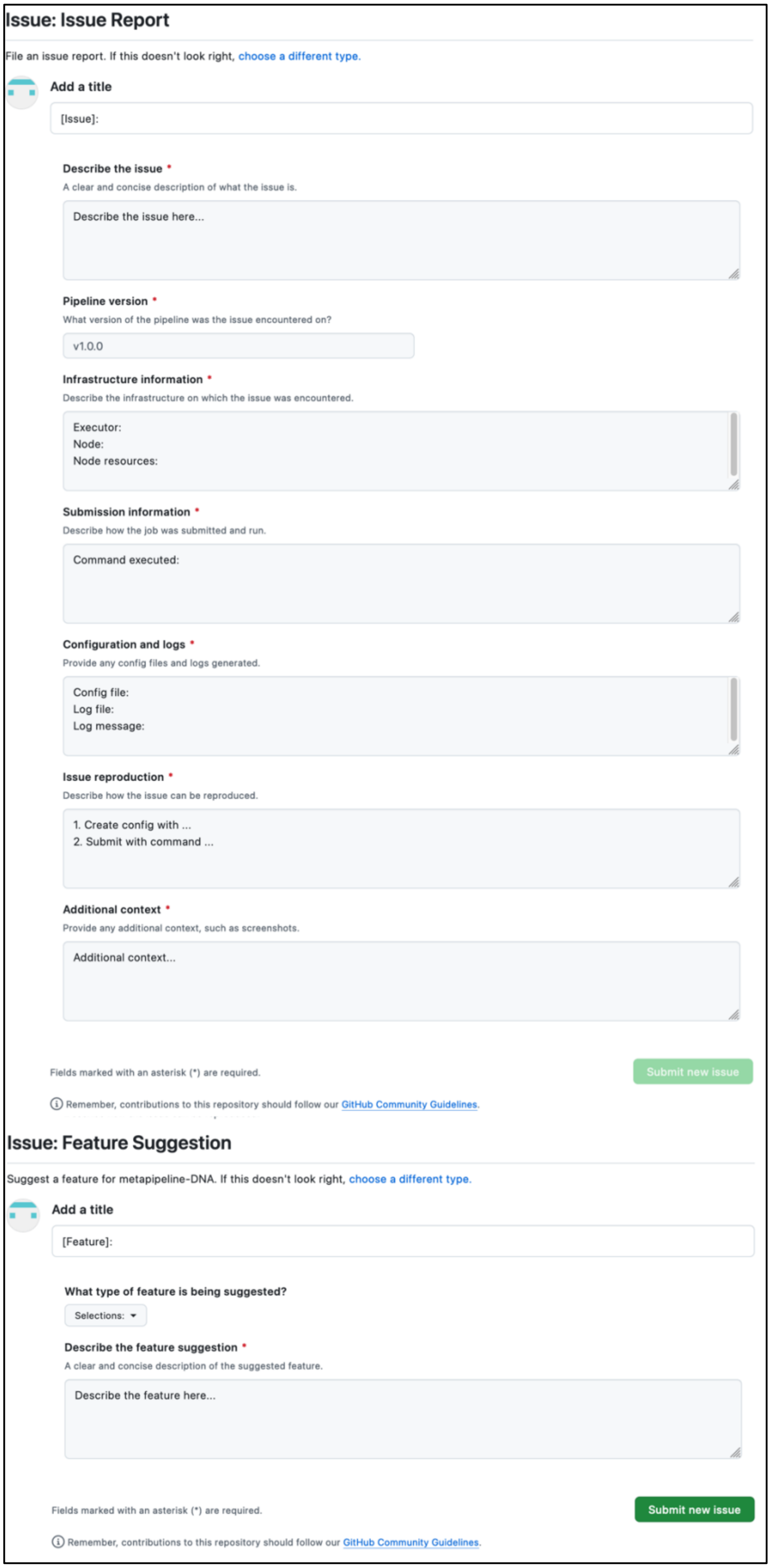
### Issue: Issue Report

File an issue report. If this doesn't look right, choose a different type.

**Add a title**

[Issue]:

**Describe the issue** *
A clear and concise description of what the issue is.

Describe the issue here...

**Pipeline version** *
What version of the pipeline was the issue encountered on?

v1.0.0

**Infrastructure information** *
Describe the infrastructure on which the issue was encountered.

Executor:
Node:
Node resources:

**Submission information** *
Describe how the job was submitted and run.

Command executed:

**Configuration and logs** *
Provide any config files and logs generated.

Config file:
Log file:
Log message:

**Issue reproduction** *
Describe how the issue can be reproduced.

1. Create config with ...
2. Submit with command ...

**Additional context** *
Provide any additional context, such as screenshots.

Additional context...

Fields marked with an asterisk (*) are required.

Submit new issue

Remember, contributions to this repository should follow our GitHub Community Guidelines.

### Issue: Feature Suggestion

Suggest a feature for metapipeline-DNA. If this doesn't look right, choose a different type.

**Add a title**

[Feature]:

**What type of feature is being suggested?**

Selections:

**Describe the feature suggestion** *
A clear and concise description of the suggested feature.

Describe the feature here...

Fields marked with an asterisk (*) are required.

Submit new issue

Remember, contributions to this repository should follow our GitHub Community Guidelines.

**Supplemental Figure 1**: **Reporting templates.** Issue forms for submitting bug reports and feature suggestions, with structured input options to describe the bug/feature.

```
.
└── metapipeline-DNA-6.0.0
    └── PROJECT-ID
        ├── log-metapipeline-DNA-timestamp
        │   ├── nextflow-log
        │   │   ├── PATIENT1-ID-workdir
        │   │   └── PATIENT2-ID-workdir
        │   └── process-log
        │       ├── align_DNA
        │       ├── call_gSNP
        │       ├── call_gSV
        │       ├── call_metapipeline_DNA
        │       ├── call_mtSNV
        │       ├── call_sCNA
        │       ├── call_SRC
        │       ├── call_sSNV
        │       ├── call_sSV
        │       ├── check_process_status
        │       ├── generate_SQC_BAM
        │       └── recalibrate_BAM
        └── main_workflow
            └── output
                ├── align-DNA-10.0.0
                ├── call-gSNP-10.0.1
                ├── call-gSV-5.2.0
                ├── call-mtSNV-4.0.0
                ├── call-sCNA-3.2.0
                ├── call-SRC-2.0.0
                ├── call-sSNV-8.1.0
                ├── call-sSV-6.1.0
                ├── generate-SQC-BAM-1.0.0
                └── recalibrate-BAM-1.0.1
                    └── SAMPLE-ID
                        ├── GATK-4.2.4.0
                        │   └── output
                        │       └── BWA-MEM2-2.2.1_GATK-4.2.4.1_DATASET-ID_SAMPLE-ID.bam
                        └── log-recalibrate-BAM-timestamp
```

**Supplemental Figure 2: Output and log directory structure**. Outputs are organized under each pipeline with a sample/patient/project identifier followed by a directory for logs and a directory for each main tool used in the pipeline. Metapipeline-DNA outputs follow the same structure with individual pipelines' outputs organized recursively in metapipeline-DNA's output.

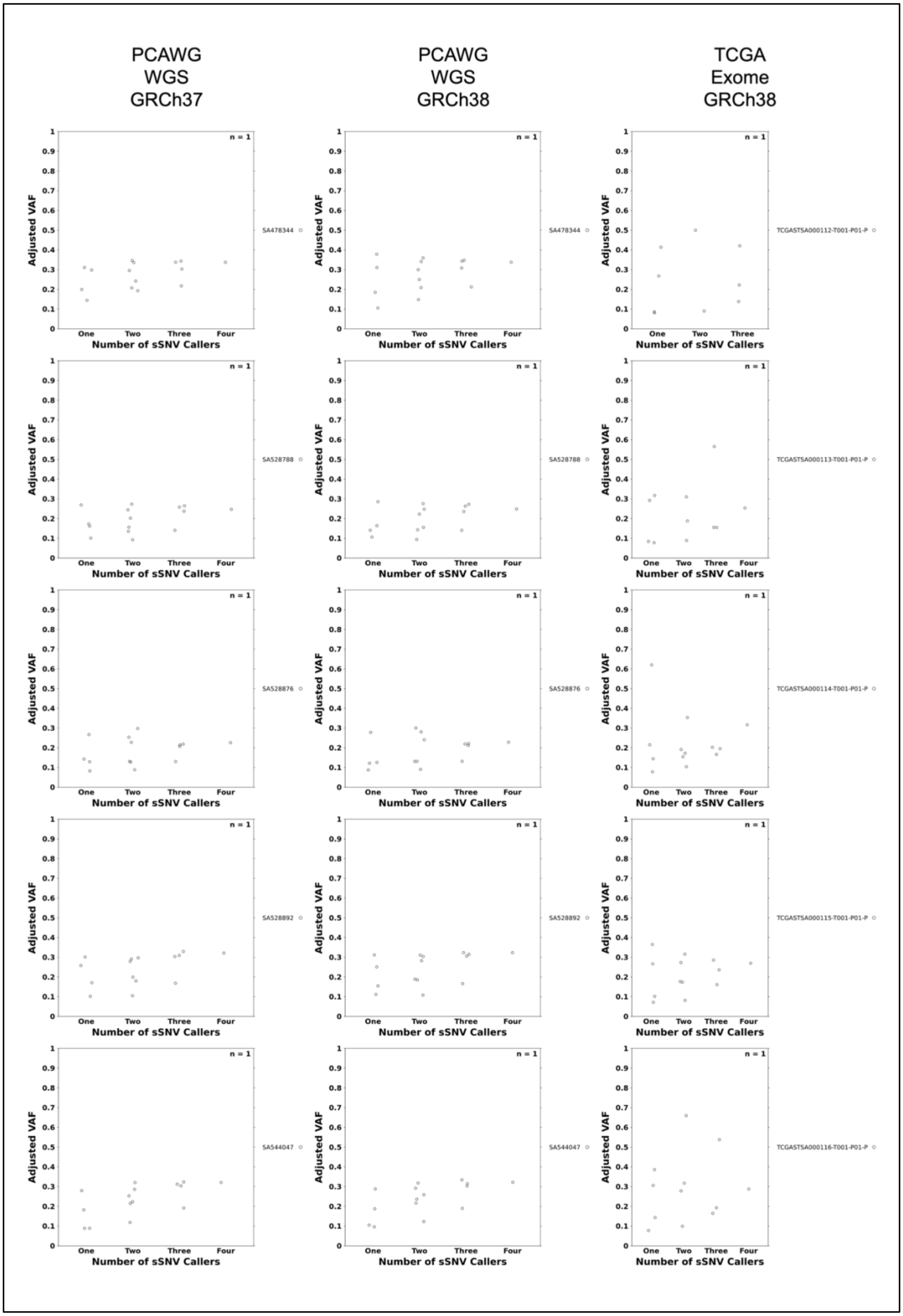


**Supplementary Figure 3: VAF plots for all samples.** Variant allele frequencies based on consensus between callers for all samples processed.
